# Versatile and multiplexed mass spectrometry-based absolute quantification with cell-free-synthesized internal standard peptides

**DOI:** 10.1101/2021.05.16.444331

**Authors:** Keiko Masuda, Keiko Kasahara, Ryohei Narumi, Masaru Shimojo, Yoshihiro Shimizu

**Author notes:** Correspondence should be addressed to: **Yoshihiro Shimizu, Ph.D.** Laboratory for Cell-free Protein Synthesis, RIKEN Center for Biosystems Dynamics, Research (BDR), 6-2-3 Furuedai, Suita, Osaka 565-0874, Japan.

## Abstract

Preparation of stable isotope-labeled internal standard peptides is crucial for mass spectrometry (MS)-based targeted proteomics. Herein, we developed versatile and multiplexed absolute protein quantification method using MS. A previously developed method based on the cell-free peptide synthesis system, termed MS-based quantification by isotope-labeled cell-free products (MS-QBiC), was improved for multiple peptide synthesis in one-pot reaction. We pluralized the quantification tags used for the quantification of synthesized peptides and thus, made it possible to use cell-free synthesized isotope-labeled peptides as mixtures for the absolute quantification. The improved multiplexed MS-QBiC method was proved to be applied to clarify ribosomal proteins stoichiometry in the ribosomal subunit, one of the largest cellular complexes. The study demonstrates that the developed method enables the preparation of several dozens and even several hundreds of internal standard peptides within a few days for quantification of multiple proteins with only a single-run of MS analysis.

## Introduction

In terms of both high sensitivity and accuracy, mass spectrometry (MS) is becoming one of the most dominant approaches for protein identification and quantification. A key technique for the protein quantification is the use of a stable isotope-labeled (SIL) peptide as an internal standard. We cannot simply quantify the amount of peptides based on their signal intensity values since the ionization efficiency of a peptide differs according to their physical properties. The quantification is achieved by utilizing a SIL peptide at a known concentration, with the identical molecular property to a target peptide. The target peptide (a light peptide) can be quantified by comparing the signal intensities of the target peptide with those of an internal standard peptide (a heavy peptide).

Various methods are available for preparation of SIL peptides. Chemical labeling such as dimethyl labeling [1] is a useful technique for comparative quantification by labeling samples with both light and heavy tags. Another type of chemical labeling, using isobaric tags, including TMT [2], iTRAQ [3], and mTRAQ [4], are commercially available and exploited for a variety of multiplexed quantification [5].

Direct synthesis of SIL peptides is also common in the quantitative proteomics field. Chemical synthesis termed AQUA [6] is the most intuitive approach for the absolute quantification. Preparation of internal standards can be achieved in a manner with high yield and high purity, without unnecessary additional sequences. The use of non-canonical amino acids in the chemical reactions enables preparation of peptides with post-translational modifications such as phosphorylation, methylation, acetylation, and amidation. Application of a cellular protein synthesis system, such as SILAC [7, 8], is suitable for the preparation of peptides or proteins that are difficult to be chemically synthesized, which is an alternative way for the direct synthesis of the SIL peptides. However, these approaches have some drawbacks such as high cost using expensive labeled amino acids and the need to control its metabolism, which result in difficulties in applying them to a large scale studies.

As an alternative and effective method for the preparation of internal standards, a cell-free protein synthesis system has been getting popular in recent years [9]. The history of the cell-free protein synthesis system is so long, dating back to a report in 1954 which showed that rat liver extract has an amino acid polymerization ability [10]. Presently, various cell-free systems are available, originating from *Escherichia coli*, archaea, protozoan, yeast, wheat germ, tobacco, insects, and mammals [11]. Compared to chemical or cellular synthesis methods, the cell-free system can be performed in micro-litter (pico-mol yield) scale, hence it is easy to minimize the use of expensive SIL amino acids.

Various forms of cell-free systems have been proposed, of which the reconstituted system, composed of only factors related to the translation and transcription systems, has some features applicable to the quantitative proteomics studies. The system, termed Protein synthesis Using Recombinant Elements (PURE) system [12], does not include nucleases and therefore, short and linear DNA templates such as PCR-amplified products could be utilized. The system also does not contain proteases nor metabolic enzymes, and therefore, unexpected degradation of synthesized peptides and stable isotope scrambling [13] could be avoidable. These features are valuable for ensuring quality of synthesized internal standards and resultantly precise quantification.

By utilizing the PURE system, we have previously established a cost-effective absolute protein quantification method, termed MS-based Quantification By isotope-labeled Cell-free products (MS-QBiC) [14]. The method uses a unique design of an internal standard peptide composed of a FLAG tag, a spacer peptide, a quantification tag, and any target peptide, which we term the MS-QBiC peptide (**Fig. 1**). The sequence of the quantification tag “LVTDLTK” is a tryptic peptide derived from bovine serum albumin (BSA) that is easily available from the public resources. Therefore, it is easy to estimate the amounts of MS-QBiC peptides by adding a known amount of BSA. After tryptic digestion of mixtures containing a sample and the MS-QBiC peptide, the ratio of light and heavy peptide is analyzed using MS. Based on the known amount of light quantification tag, the amount of the target peptide derived from the sample can be quantified. Since this quantification scheme is based on the single intermediary quantification tag (LVTDLTK), there is a difficulty to apply the MS-QBiC method for the multiplexed quantification. Such a drawback can be improved by increasing the variation of the quantification tag.

**Fig. 1.**
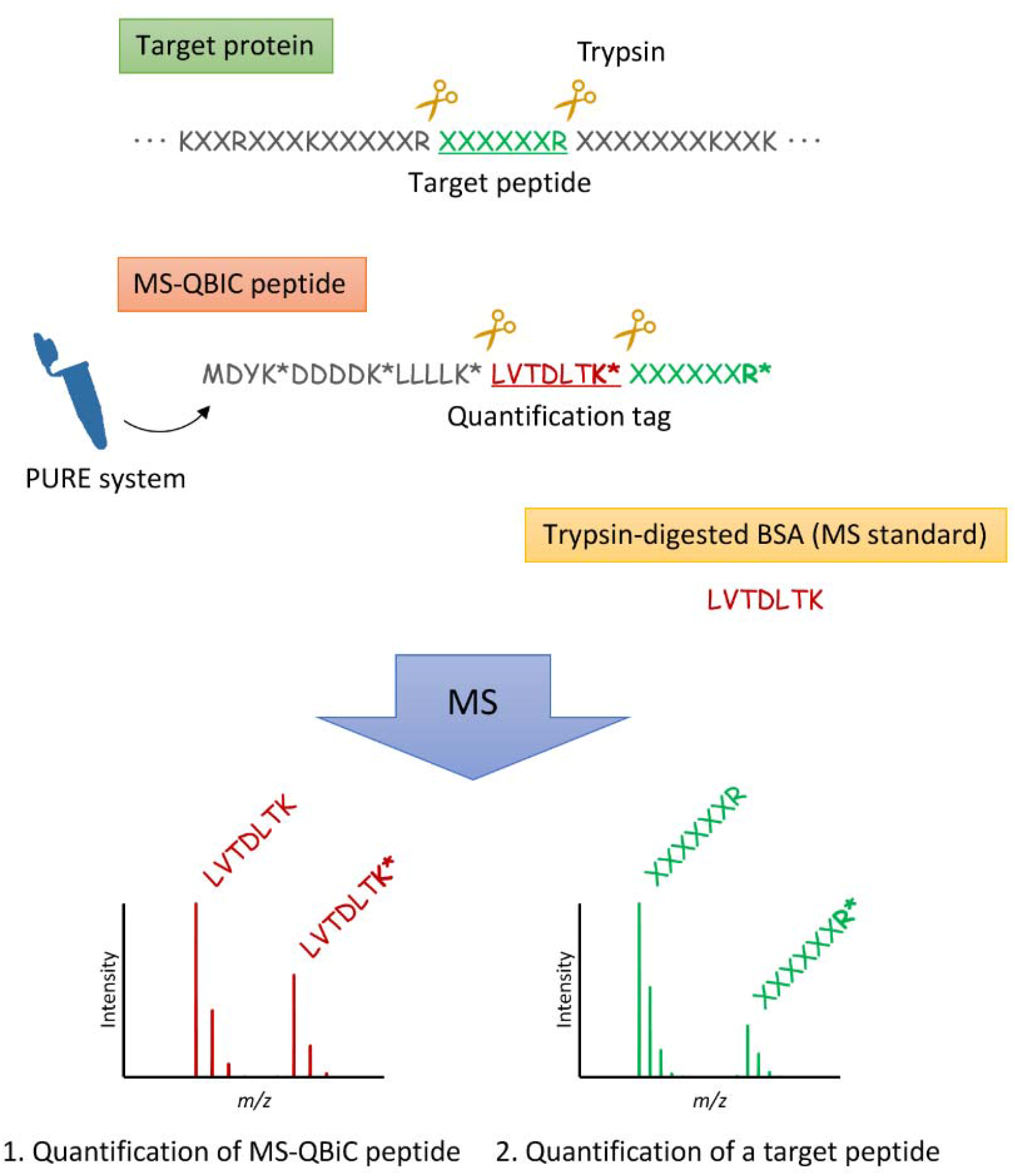
MS-QBiC method. Tryptic digestion of the MS-QBiC peptide, synthesized with the PURE system, generates a quantification tag (red) and a target peptide (green) containing stable isotope labeled arginine (R*) or lysine (K*). The target peptide derived from a target protein by tryptic digestion is analyzed with MS by comparing peak intensities of light and heavy peptides. The quantification tag is analyzed with MS by comparing peak intensities of light (derived from BSA or chemically synthesized) and heavy peptides to obtain the amount of synthesized MS-QBiC peptide.

In the present study, we improved the MS-QBiC method by designing new quantification tags. First, we designed more than 100 candidates of LVXXLTK, where X is variable amino acid, and picked up 34 tags which showed good solubility and signal intensity on MS analyses. Then, we evaluated the new 34 tags and demonstrated multiplexed MS-QBiC method using two samples. One is the commercially available tryptic peptides mixture derived from six proteins in which the quality is guaranteed. Another is the 30S ribosomal subunit from *Escherichia coli*, one of the largest cellular complexes composed of one RNA (16S rRNA) and 21 ribosomal proteins. We demonstrate that all quantified amount of proteins were within the same order of magnitude as the theoretical values, indicating usefulness of the developed method for multiplexed quantitative proteomics studies.

## Results

### Selection of new quantification tags

Based on the original quantification tag (LVTDLTK), we made a list of 110 candidates of single and double point mutants, LVXXLTK, by replacing third Thr and fourth Asp with 11 amino acids, including Ala, Asp, Glu, Phe, Gly, Leu, Pro, Ser, Thr, Val, and Tyr (**Table S1**). We did not use other amino acids for variable reasons. Lys and Arg were excluded to avoid internal tryptic digestion. Met, Cys, and Trp were not used because they can be oxidized. Asn and Gln were amino acids potentially be deamidated. We also avoided His that tends to show week MS intensity [15]. Ile was excluded because it is a structural isomer of Leu.

Resultant 110 candidates were arranged in the order of *m/z* (*z* = 2) and the number was reduced to 42 to keep proper intervals at least more than 0.2 *m/z* (**Table S1**). The selection was performed to avoid extracted-ion chromatograms (XICs) become complicated. We note that *m/z* of ^13^C_6_ Lys labeled peptides (heavy peptides) were also considered in this selection scheme. Next, the selected 42 candidates were chemically synthesized by Fluorenylmethyloxycarbonyl (Fmoc) solid-phase peptide synthesis. Hydrophobic peptides tended to be problematic, where some peptides, such as LVLLLTK, LVLFLTK, and LVFFLTK, resulted in low yield and some peptides, such as LVVLLTK, LVVFLTK, LVVYLTK, LVYLLTK, and LVFYLTK, resulted in insoluble in water. After the selection based on the characteristics of the actually synthesized peptides, we obtained 34 quantification tags (**Table S2**).

Using a series of chemically synthesized peptides, we prepared a quantification tag mixture containing 100 µM each of 35 peptides including original LVTDLTK and newly designed 34 tags. Concentration of each peptide was measured by amino group determination [16]. According to the designed 34 tags, we also prepared 34 plasmids encoding each new quantification tag by site-directed mutagenesis using the original plasmid encoding LVTDLTK.

### Multiplexed cell-free peptide synthesis and quantification

To validate the newly designed tags, we tried to measure protein mixtures with known concentration using the MS-QBiC method based on 34 quantification tags. As a protein mixture, we used a commercially available equimolar tryptic peptide mixture from six proteins including bovine cytochrome C (12 kDa), chicken lysozyme C (16 kDa), yeast alcohol dehydrogenase 1 (37 kDa), bovine serum albumin (69 kDa), bovine serotransferrin (78 kDa), and *E. coli* beta-galactosidase (117 kDa) (Pierce 6 Protein Digest, equimolar, Thermo Scientific). A total of 34 target peptides was selected for 6 proteins quantification (**Data S1** and **Table S3**). For the selection of the target peptides, we focused on those composed of 6 to 20 amino acids without Met and Cys. Also, the peptide sequences that may cause miss-cleavages, such as KP, RP, KK, and RR were avoided. We note that the original quantification tag, LVTDLTK, was not used for this study because it was derived from bovine serum albumin, a component of the target protein mixture.

A mixture of 34 MS-QBiC peptides were synthesized in a single PURE system reaction. DNA templates for each peptide synthesis were amplified in separate PCR reactions, and then, all 34 templates were added to the PURE system and peptides were synthesized in a one pot reaction. The resultant mixture of 34 MS-QBiC peptides was mixed with chemically synthesized non-labeled quantification tags and six proteins mixture (total 1 pmol, 167 fmol each) and then further processed for MS analysis including tryptic digestion.

The yield of each MS-QBiC peptide was measured by LC-MS analysis using Orbitrap mass spectrometer in full-scan mode (**Fig. 2a**). The measurement was based on the light/heavy ratio of the quantification tags and it showed that all 34 MS-QBiC peptides were successfully synthesized. The yields from 5 µl PURE reaction mixture ranged between 11 and 126 fmol and the total amounts were 2 pmol. They were within the range for the MS-based quantification, although some peptides, such as YVVDTSK*, AWSVAR*, LVNELTEFAK*, YYGYTGAFR*, and LWSAEIPNLYR*, showed low yield (asterisks represent labeled lysine or arginine). We note that additional synthesis of peptides can compensate for the low yield of specific peptides. When YVVDTSK*, LVNELTEFAK*, and LWSAEIPNLYR* were synthesized separately, the yield from 1 µl of the PURE reaction were 200, 270, and 120 fmol, respectively. Therefore, supplementation of the separately synthesized peptides can be performed if necessary.

**Fig. 2.**
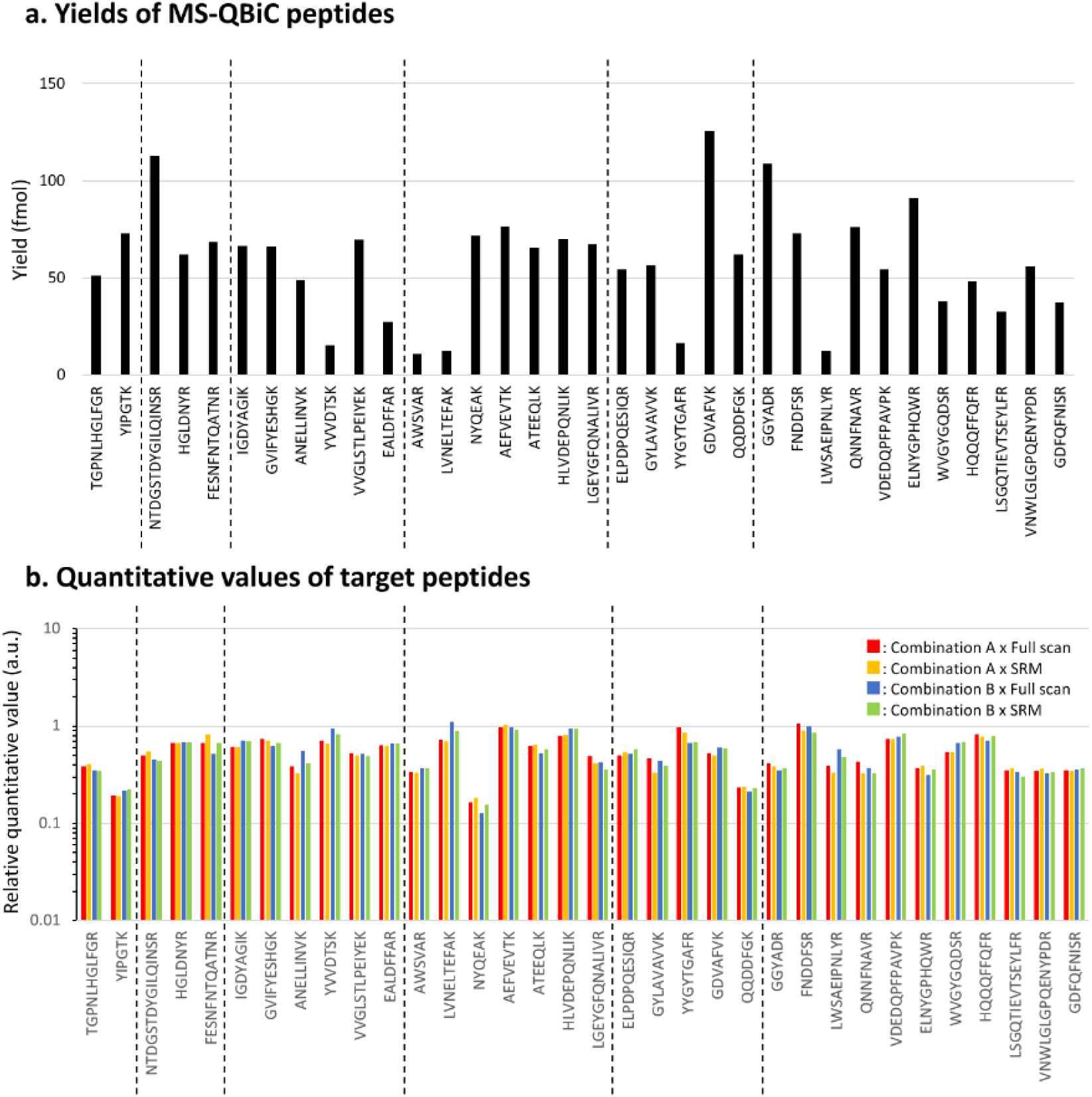
Multiplexed MS-QBiC method using newly designed 34 quantification tags. (a) Yields of MS-QBiC peptides. A total of 34 MS-QBiC peptides were synthesized in a single PURE reaction and the yields of each peptide were quantified according to the light/heavy ratio of the quantification tags. Yields from 5 μL PURE reaction mixture are shown. (b) Quantitative values of target peptides. Ratios of the calculated quantitative value to the input protein amount are shown as relative quantitative values. The results obtained with LC-MS analysis (red and blue) and SRM analysis (yellow and green) using MS-QBiC peptides in combination A (**Table S4**) (red and yellow) and those using combination B (blue and green) are shown.

Subsequently, target peptides in a six proteins mixture were quantified based on the light/heavy ratio of the target peptides by LC-MS analysis (**Fig. 2b**, red). We also performed selected reaction monitoring (SRM) using triple quadrupole mass spectrometer and found that the quantification results were almost similar with those with LC-MS analysis (**Fig. 2b**, orange). Further validation was performed by shuffling the combination of the quantification tags and target peptides by reversing the order of the quantification tags (**Table S4**) and target peptides were quantified with both LC-MS and SRM. We found that the quantification results did not vary according to the used quantification tags (**Fig. 2b**, blue and green).

It was possible that quantitative values of 34 target peptides can vary due to various factors such as miss-cleavage, non-specific modification, hydrophobicity, and detectability. Detailed analysis of the ion chromatograms showed the miss-cleavage product of YIPGTK (**Fig. S1a**). Also deamidated products originated from NYQEAK and QQDDFGK were found (**Fig. S1b, c**). NYQEAK also showed early retention time, suggesting it is very hydrophilic and not sufficiently retained by the column (**Fig. S1b)**. As a result, these peptides were quantified at relatively low values (**Fig. 2b**). After excluding the values of these three peptides, average of quantification values of cytochrome C, lysozyme C, alcohol dehydrogenase 1, serum albumin, serotransferrin, and beta-galactosidase were 62, 102, 103, 113, 95, and 86 fmol, respectively, which were within the same order of magnitude as the amounts of added proteins (167 fmol).

### Quantification of ribosomal 30S subunit

Further verification of the developed method was performed by analyzing the ribosomal proteins stoichiometry in *E. coli* ribosomal 30S subunit composed of 21 ribosomal proteins. Quantifying ribosome composition is crucial for understanding its biogenesis [17, 18]. Also it might be important because the ribosome stoichiometry is suggested to be variable depending on tissue types, physiological conditions, and aging, which might cause phenotypic changes in eukaryotes [19–21].

A total of 68 peptides were designed (**Data S2** and **Table S5**) and two sets of the PURE reaction were performed where 34 peptides were synthesized in each reaction. Synthesized peptides were quantified with chemically synthesized non-labeled quantification tags and then mixed as a solution for 30S subunit quantification containing 68 MS-QBiC peptides.

*E. coli* ribosomal 30S subunit (100 fmol), in which the concentration was determined with UV absorbance, was mixed with the cell-free synthesized peptide solution and each peptide was quantified. As with the result of six proteins quantification, most of the peptides were within the same order of magnitude as the amount of added 30S subunit (**Fig. 3a**). Five peptides marked with asterisks (ISELSEGQIDTLR from uS13, AIISDVNASDEDR from uS14, WNAVLK from uS14, GPFIDLHLLK from S19, and ENEPFDVALR from S21) were less than 30% compared to the added 30S subunit amount. Detailed analysis of the ion chromatograms again showed the miss-cleavage of these peptides (**Fig. S2**), suggesting these peptides are not suitable for the quantification. After excluding the values of these five peptides, protein amounts were calculated as the average of quantification values of individual peptides (**Fig. 3b**). The data showed that all proteins were within the same orders of magnitude, suggesting equimolar amount of proteins are included in the prepared 30S subunit.

**Fig. 3.**
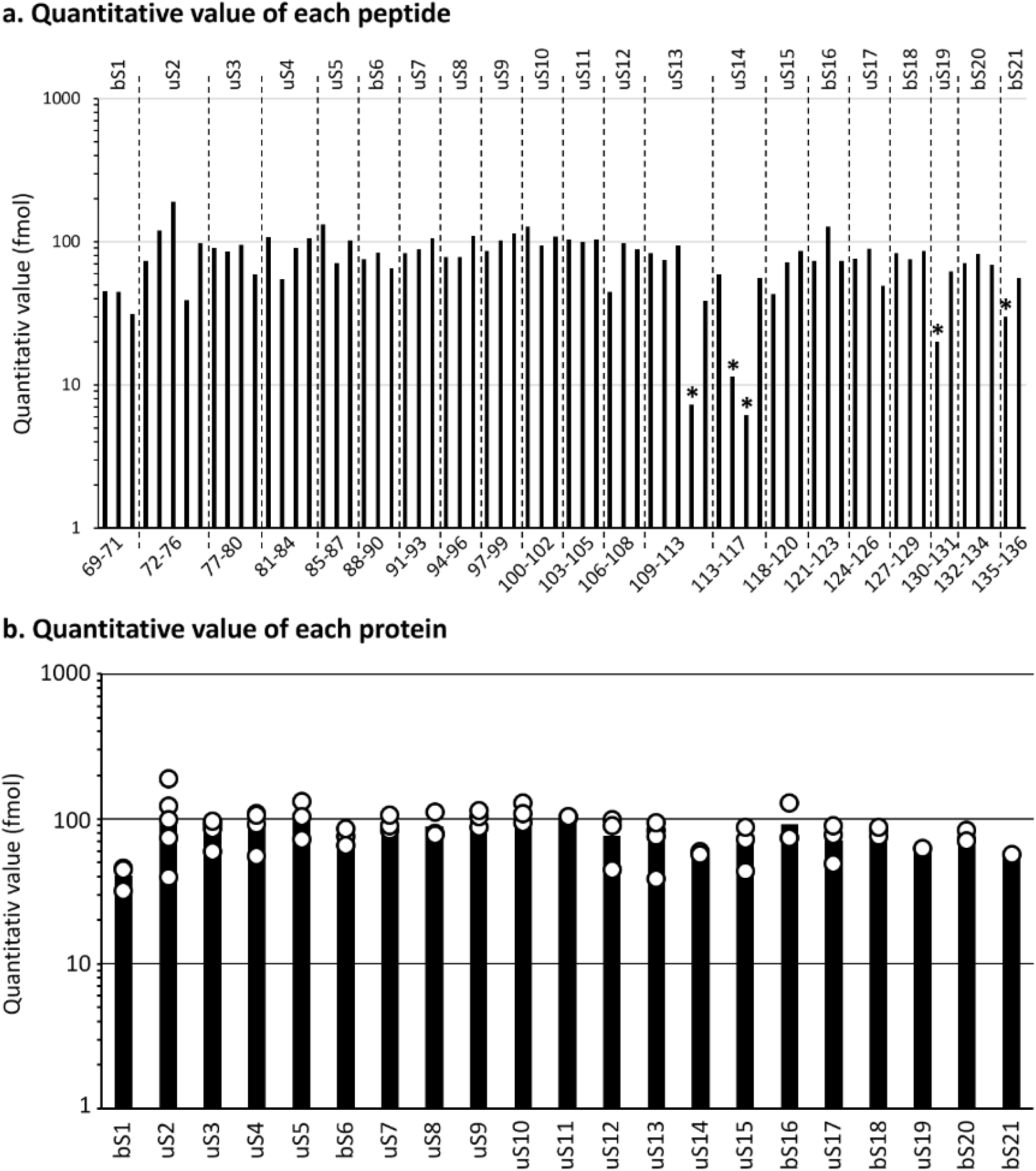
Quantification of ribosomal 30S subunit. (a) Quantitative values of each peptide. Target peptides derived from 21 ribosomal proteins in 100 fmol ribosomal 30S subunit was quantified with 68 MS-QBiC peptides. Asterisks represent peptides less han 30% compared to the added 30S subunit. (b) Quantitative values of each protein. Protein amounts calculated as the average of quantitative values of individual peptides are shown. The peptides marked with asterisks in (a) were not included in the calculation because of the miss-cleavage issues. Each dot represents the quantitative value of individual peptides.

## Discussion

Mass spectrometric methods have outstanding advantages in sensitivity and accuracy for protein quantification. However, a variety of preliminary surveys are required for analyzing the samples of interest. In order to obtain an accurate quantification result, it is indispensable to select appropriate target peptides, which are effectively ionized and detectable on MS (“proteotypic”) and quantitatively reliable (“quantotypic”) [22, 23]. Quantotypic peptides are those that do not have chemical and/or post-translational modification sites and also do not have sequences which are susceptible to incomplete tryptic digestion. Generally, screening of quantotypic peptides is performed computationally and experimentally that takes much time and effort [24, 25], and then, their SIL counterparts are synthesized. When SRM method is applied, further practical screening of reliable transitions of target peptides are necessary. There could be a case that preparing all target peptides as a SIL form is difficult for financial reasons. Furthermore, there is a problem that the practical screening is not always applicable when acquiring standard proteins are difficult.

By using the approach presented here, it is possible to prepare internal standard peptides without limiting the number of peptides to be synthesized, which may result in more practical screening of quantotypic peptides. By increasing the variation of the quantification tag, the throughput of both the internal peptides preparation and the sample quantification can be highly improved because multiplexed quantification tags make it possible to synthesize at most 35 peptides in one pot reaction. Preparation steps including PCR amplification, cell-free peptide synthesis, FLAG affinity purification, and tryptic digestion are finished only within two days.

Two quantification steps are required to quantify the samples of interest, in that synthesized MS-QBiC peptides are quantified first and then the samples are quantified with the MS-QBiC peptides as references. It is noteworthy that these multi-step quantification can be completed by a single run of LC-MS analysis. We used the orbitrap mass analyzer at full scan mode which covers whole information of target peptides and miss-cleaved peptides. Target peaks are easily identified by contrasting light peptides and heavy peptides eluting at the same retention time, not even need MS/MS characterization. Kumar *et al*. called such an approach, MS western, originating from a western blotting [26]. MS/MS-free method does not waste time acquiring MS2 and therefore enables short and high-throughput analysis [27]. After processing the peak data and calculation, some peptides show outliers due to miss-cleavages, noisy peaks, or missing peaks. Those discordances can be corroborated by analyzing back the full scan data and the outliers will be eliminated accordingly. Whole processes will be completed within a weekday: Designing and acquisition of reverse primers encoding target peptides (2 days including a shipping time), PCR amplification of DNA template for the PURE system (3 hours, for the first time only), synthesis of SIL peptides using the PURE system (2 hours including a purification time), tryptic digestion of samples and MS-QBiC peptides (overnight), and MS analysis (a few hours per sample) (**Figure 4**).

**Fig. 4.**
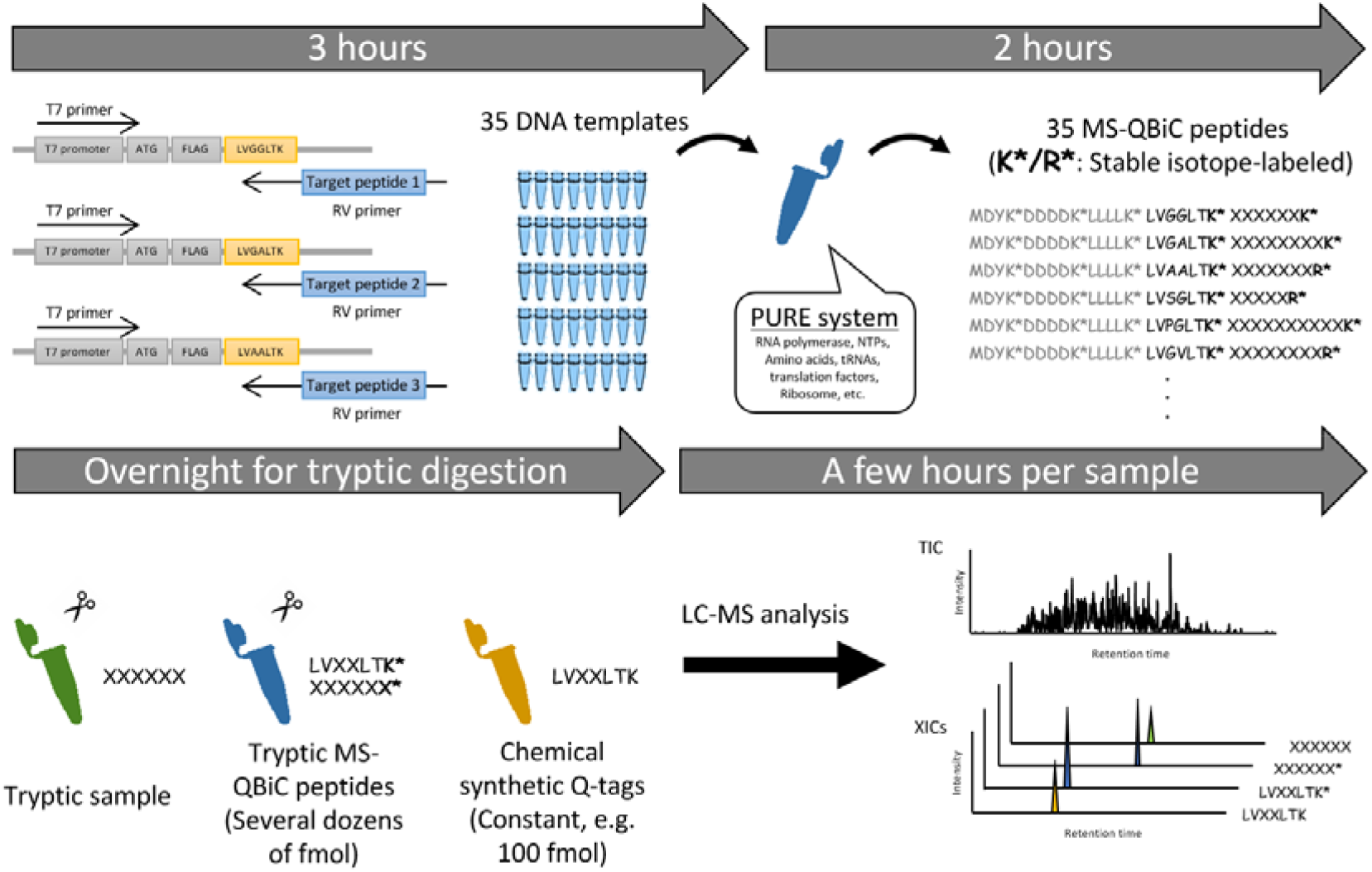
Workflow of multiplexed MS-QBiC method. A total of 35 MS-QBiC peptides can be obtained in 5 hours using only one PURE reaction mixture. After tryptic digestion with sample proteins overnight, quantification results based on the LC-MS analysis can be obtained in the next day.

By establishing new 34 quantification tags, now we can basically perform the 35-plex quantification by a single run of MS analysis including both steps of quantification of cell-free synthetic heavy peptides and quantification of target peptides in samples. Furthermore, we showed that it is possible to increase the multiplexity by using multiple sets of MS-QBiC peptides if the quantification of the MS-QBiC peptides and samples are separated. For the quantification of the ribosomal proteins in 30S subunit, we designed and prepared 68 peptides as two sets of 34 MS-QBiC peptides (**Table S6, Data S2**). Then, in the first step, each set of MS-QBiC peptides was quantified with chemically synthesized quantification tags. Next, two sets of 34 MS-QBiC peptides were mixed and 68 MS-QBiC peptides mixture was obtained. Finally, the mixture containing known amount of 68 MS-QBiC peptides was spiked into the sample and analyzed to quantify the endogenous target peptides. Using this approach, increasing the multiplexity is possible for quantification of the multi-protein complex such as the ribosome as shown here and/or measuring a systems-level dynamics of biological systems, *e*.*g*., signal transduction networks.

## Materials and Methods

### Preparation of plasmids encoding quantification tags

Preparation of plasmid encoding an original quantification tag (LVTDLTK) was already reported [14]. Briefly, a DNA sequence (5′-ATGGACTACAAGGACGACGACGACAAGCTGCTGCTGCTGAAGCTGGTTACTGACCTGACTAAG-3′) that encodes tandemly arranged FLAG tag, spacer peptide, and a quantification tag (MDYKDDDDK-LLLLK-LVTDLTK) was cloned into a pURE1 vector (BioComber) containing a T7 promoter sequence and the ribosome-binding site. Using the original plasmid as a template, 34 plasmids encoding new quantification tag (LVXXLTK) were amplified using PfuUltra II Fusion HS DNA Polymerase (Agilent Technologies) using appropriate DNA primers (**Table S7**). After amplification, restriction enzyme Dpn I (Takara Bio) was added to cut and remove the original template plasmid at 37 °C overnight. Using the remaining PCR products, *E. coli* JM109 competent cells (Toyobo) were transformed and cultured on LB agar plate containing ampicillin at 37 °C overnight. Formed colonies were picked up and re-cultured in LB medium containing ampicillin at 37 °C overnight. The resultant plasmids were purified using QIAprep Spin Miniprep Kit (Qiagen).

### Preparation of non-labeled peptides by chemical synthesis

Non-labeled quantification tags were synthesized by Fluorenylmethyloxycarbonyl (Fmoc) chemistry using a peptide synthesizer SyroWave (Biotage, Uppsala, Sweden). Fmoc amino acids were purchased from Watanabe Chemical Industries and contained the following side chain protecting groups: Asp(OtBu), Glu(OtBu), Ser(tBu), Thr(tBu), and Tyr(tBu). Fmoc-Lys(Boc)-TrtA-PEG resin (loading 0.21 mmol/g), 1-[Bis(dimethylamino)methyliumyl]-1H-benzotriazole-3-oxide hexa fluorophosphate (HBTU), and *N*,*N-*diisopropylethylamine (DIEA) were also obtained from Watanabe Chemical Industries. Piperidine was purchased from Nacalai Tesque (Kyoto, Japan). Fmoc deprotection was performed using 40% piperidine/DMF for 3 + 12 min. Coupling was performed using Fmoc amino acids/HBTU/DIPEA (equivalents = 5:5:10) for 60 min. After the synthesis, the products were washed with dichloromethane and dried. The peptides were cleaved from the resin with reagent K (82.5% TFA, 5% phenol, 5% thioanisol, 5% H_2_O, and 2.5% 1,2-ethanedithiol) for 3 h at room temperature. Cleaved peptides were precipitated and washed with pre-cooled diethyl ether, and dried. The product was purified by reverse phase high-performance liquid chromatography (RP-HPLC) using a YMC-Pack Pro C18 column (10 × 150 mm; YMC, Kyoto, Japan) with a linear gradient of acetonitrile containing 0.1% TFA with a flow rate of 1.0 ml/min. The elution was monitored at 220 nm. Collected peak fraction was evaporated and lyophilized.

Lyophilized peptides were dissolved in 2% acetonitrile containing 0.1% TFA. The concentrations were measured by an amino group determination method [16]. Then, aliquot of them were mixed to make a stock mixture at a concentration of 100 µM and stored at −80 °C until use.

### Preparation of heavy-labeled peptides by cell-free synthesis

Using the plasmids encoding each quantification tag as templates, DNA templates for MS-QBiC peptide synthesis were amplified by PCR using Taq DNA polymerase (NEB) with a T7 promoter primer (5′-GGGCCTAATACGACTCACTATAG-3′) as a forward primer and appropriate reverse primers listed in **Table S4 and Table S6**. PCR product mixtures (2.5 μL) were directly added to yield 50 μL PURE system reaction mixtures [28] containing ^13^C_6_ ^15^N_4_ L-Arginine (R*) and ^13^C_6_ L-Lysine (K*) (Thermo Scientific) as a substitute for non-labeled L-Arginine and L-Lysine, respectively. The mixtures were incubated at 37 °C for 60 min. The synthesized peptides were purified using 10 μL slurry of anti-FLAG M2 Magnetic Beads (Sigma Aldrich) according to the manufacturer’s instruction. The peptides were eluted with 20 μL of 0.1% TFA and dried with SpeedVac.

### Reduction, alkylation, and tryptic digestion

Enzymatic digestion was performed basically according to a phase transfer surfactant (PTS)-aided protocol [29]. MS-QBiC peptides or ribosomal 30S subunit, prepared according to the previous report [18], were dissolved in 10 µl of PTS buffer (10 mM sodium deoxycholate, 10 mM sodium N-lauroylsarcosinate, and 50 mM NH_4_HCO_3_), reduced with 10 mM TCEP at 37 °C for 30 min, alkylated with 20 mM iodoacetamide at 37 °C for 30 min, and quenched with 20 mM L-cysteine residues. Digestion was performed by adding 100 ng of trypsin (Thermo Scientific) and 100 ng of Lys-C (Thermo Scientific) at 37 °C overnight. After the digestion, 1 µl of 10% TFA was added to the sample to precipitate the detergents. After the sample was centrifuged at 15,000 × g for 5 min at 4 °C, supernatant was desalted by using self-prepared stage tips [30] and dried with SpeedVac. The dried peptides were dissolved in 5% acetonitrile containing 0.1% TFA and stored at −80 °C until use. Commercially available tryptic peptides mixture (Pierce 6 Protein Digest, equimolar, Thermo Scientific) was just dissolved in H_2_O at a final concentration of 1 pmol/µl and stored at −80 °C until use.

### Mass spectrometric analysis

Two types of mass spectrometers were used. One was an Orbitrap mass spectrometer (positive mode, scan range of 200–1,500 m/z, 60,000 FWHM resolution, LTQ Orbitrap Velos Pro, Thermo Scientific), which was used for a high-resolution full-scan analysis. The other was a triple quadrupole mass spectrometer (positive mode, Q1 and Q3 resolutions of 0.7 FWHM, a cycle time of 1 sec, a gas pressure of 1.0 mTorr, TSQ Vantage, Thermo Scientific), which was used for SRM analyses. Both of them were equipped with a nanospray ion source (Nanospray Flex, Thermo Scientific) and a nano-LC system (UltiMate 3000, Thermo Scientific). Peptides were concentrated using a trap column (0.075 × 20 mm, 3 µm, Acclaim PepMap 100 C18, Thermo Scientific) and then separated using a nano capillary column (0.1 × 150 mm, 3 µm, C18, Nikkyo Technos) at a flow rate of 500 nL/min using two mobile phases A (0.1% formic acid) and B (acetonitrile and 0.1% formic acid). For the analysis of tryptic six protein mix, we used a gradient of 5% B for 5 min, 5–35% B in 65 min, 35–90% B in 1 min, and 90% B in 4 min. For the analysis of tryptic ribosome 30S subunit, we used a gradient of 5% B for 5 min, 5–45% B in 40 min, 45–90% B in 1 min, and 90% B in 4 min. Elution was directly electrosprayed (2.2 kV) into the MS. Data analysis was performed using Skyline (v4.2.0.18305) (MacCoss Lab Software) [31]. For the full scan analyses, peak area was calculated with setting MS1 filter to a count of three (M, M+1, and M+2). For the SRM analysis, firstly primary SRM transitions and collision energies (CE) was calculated using Skyline. In principle, *m/z* values of doubly charged precursor ions and *m/z* values of singly charged fragment ions which are greater than precursor ions were used. Then, optimal SRM transitions were developed by LC-SRM analysis of tryptic MS-QBiC peptide using primary SRM transitions and by screening of transitions with strong intensities. A total of 226 SRM transitions (452 transitions for light and heavy peptides) was obtained for 34 quantification tags and 34 target peptides (**Table S8**).

## CRediT authorship contribution statement

Conceptualization: KM, KK, RN, and YS; Methodology: KM; Investigation: KM Resources: KM, KK, RN, and MS; Writing – Original Draft: KM; Writing – review & editing: YS; Visualization: KM and YS; Supervision: YS; Funding acquisition: KM and YS.

## Declaration of Competing Interest

The authors declare no conflict of interest.

## Acknowledgements

This work was supported by a Grant-in-Aid (18J01791 to KM, 17H05680 to YS) from the Japan Society for the Promotion of Science (JSPS), the Human Frontier Science Program (RGP0043/2017 to YS), the Astrobiology Center Project of the National Institutes of Natural Sciences (AB311005 to YS), CREST (JPMJCR20S4 to YS) from Japan Science and Technology Agency (JST), and an intramural Grant-in-Aid from the RIKEN Center for Biosystems Dynamics Research (to YS).

## Supplementary Figures

**Fig. S1.**
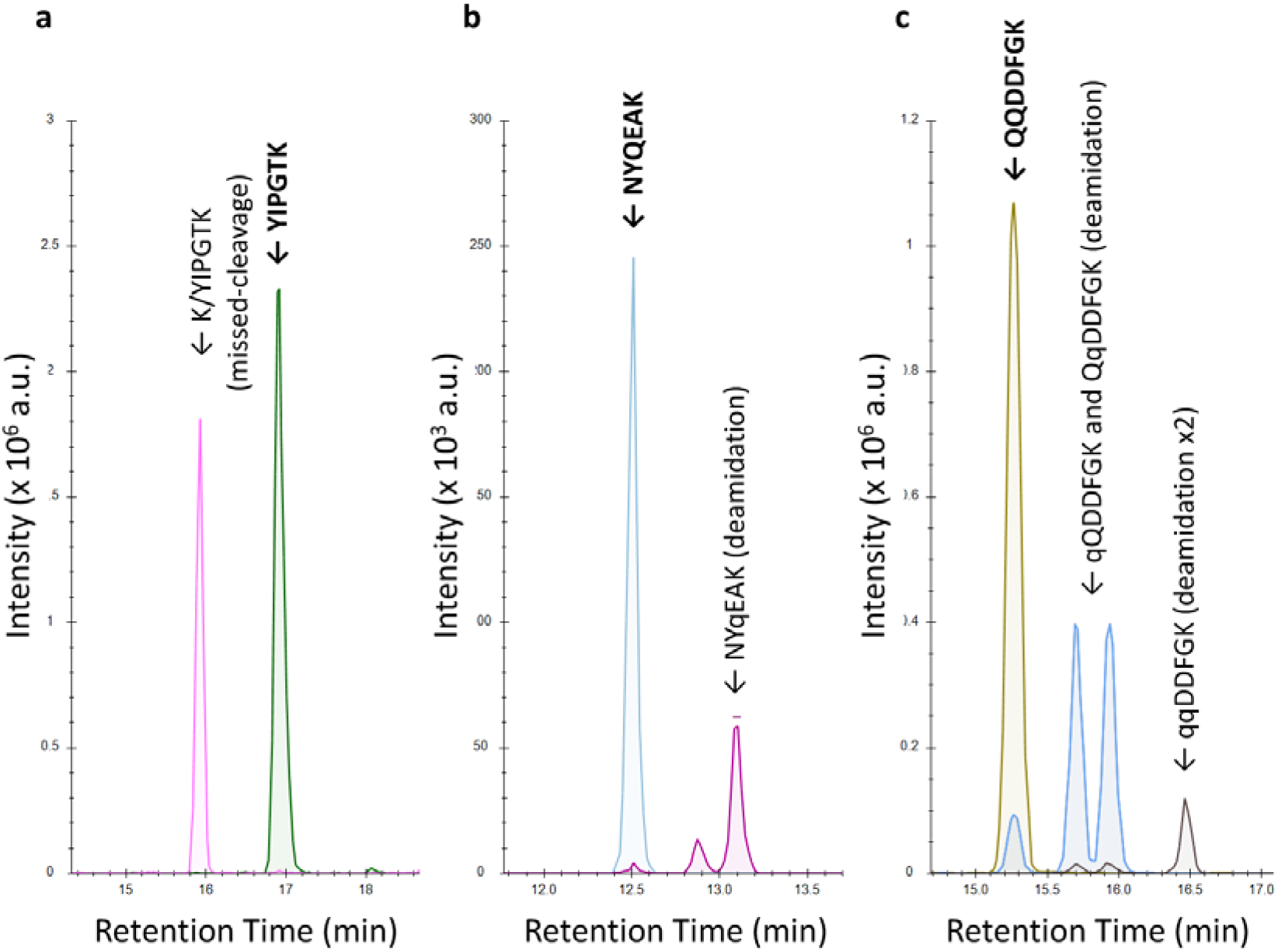
Extracted ion chromatograms of peptides problematic for quantification of six test proteins. Extracted ion chromatograms of (a) KYPGTK and YPGTK, (b) NYQEAK and NYqEAK (q represents deamidated glutamine), and (c) QQDDFGK, qQDDFGK/QqDDFGK, and qqDDFGK are shown using Skyline software [31].

**Fig. S2.**
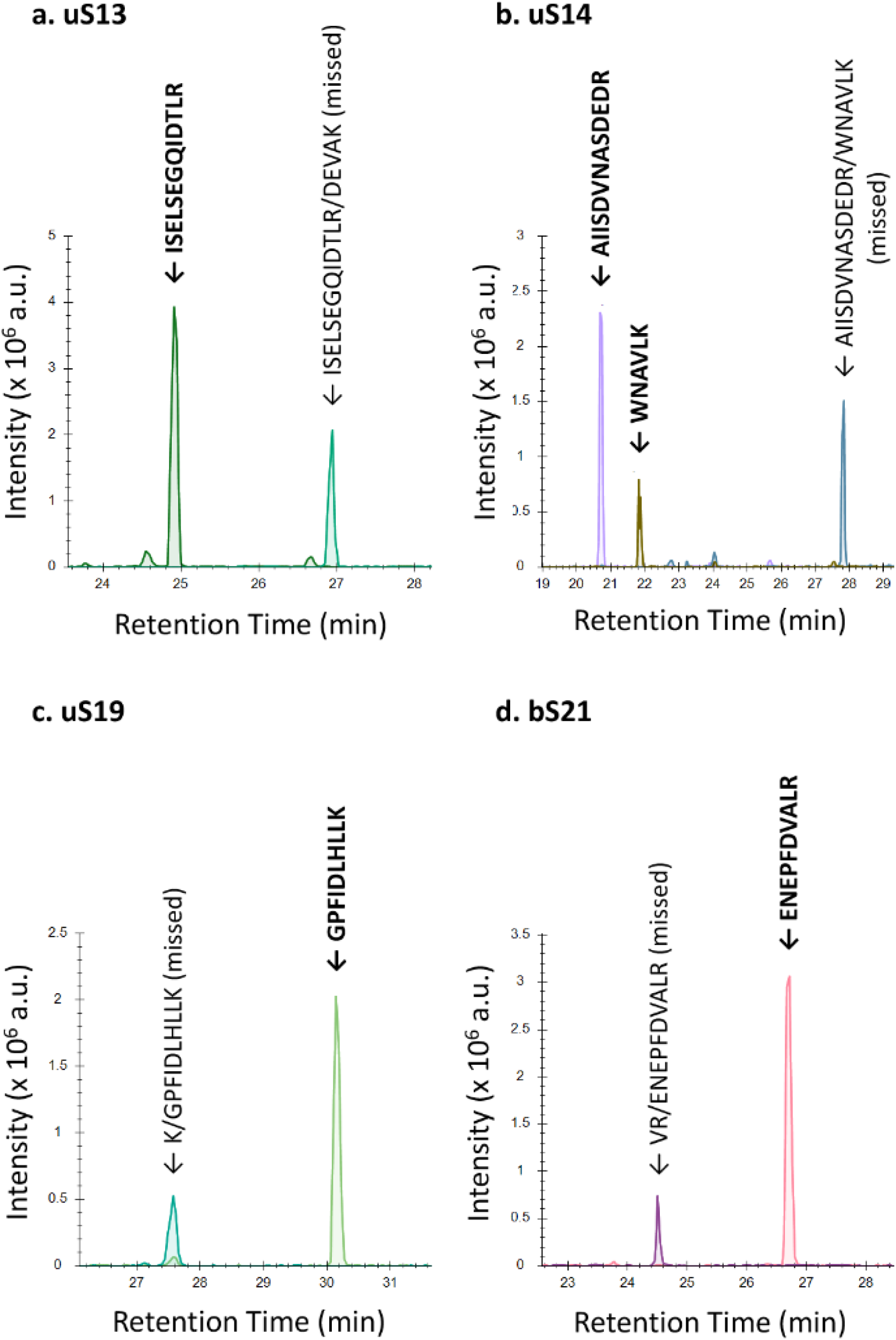
Extracted ion chromatograms of peptides problematic for quantification of ribosomal proteins. Extracted ion chromatograms of (a) ISELSEGQIDTLR and ISELSEGQIDTLRDEVAK from uS13, (b) AIISDVNASDEDR, WNAVLK, and AIISDVNASDEDRWNAVLK from uS14, (c) KGPFIDLHLLK and GPFIDLHLLK from uS19, and VRENEPFDVALR and ENEPFDVALR from bS21 are shown using Skyline software [31].

## Supplementary Tables

**Table S1. A list of 110 candidates of single and double point mutants (LVXXLTK) of original quantification tag (LVTDLTK)**. A total of 110 candidates and the original tag are listed in ascending order according to their *m/z*.

**Table S2. A list of 34 newly designed quantification tags**. A total of 34 tags are listed in ascending order according to their *m/z*.

**Table S3. A list of target peptides for quantification of six test proteins**. Designed target peptides for quantification of bovine cytochrome C, chicken lysozyme C, yeast alcohol dehydrogenase 1, bovine serum albumin, bovine serotransferrin, and *E. coli* beta-galactosidase included in Pierce 6 Protein Digest, equimolar purchased from Thermo Scientific (**Data S1**) are listed.

**Table S4. A corresponding table of quantification tags and target peptides for six test proteins**. Two combinations of quantification tags and target peptides, as well as reverse primers to synthesize corresponding peptides in a cell-free reaction, are listed.

**Table S5. A list of target peptides for quantification of 21 ribosomal proteins**. Designed target peptides for quantification of ribosomal proteins of *E. coli* ribosomal 30S subunit (bS1-bS21) (**Data S2**) are listed.

**Table S6. A corresponding table of quantification tags and target peptides for ribosomal proteins**. A combination of quantification tags and target peptides, as well as reverse primers to synthesize corresponding peptides in a cell-free reaction, are listed.

**Table S7. A list of primer sets for preparation of plasmids encoding newly designed quantification tags**.

**Table S8. A list of SRM transitions of peptides analyzed in this study**.

## Supplementary Data

**Data S1. Amino acid sequences of six test proteins**. Amino acid sequences of bovine cytochrome C, chicken lysozyme C, yeast alcohol dehydrogenase 1, bovine serum albumin, bovine serotransferrin, and *E. coli* beta-galactosidase included in Pierce 6 Protein Digest, equimolar purchased from Thermo Scientific are shown. Target peptides are shown with underlined bold characters.

**Data S2. Amino acid sequences of 21 ribosomal proteins**. Amino acid sequences of ribosomal proteins of *E. coli* ribosomal 30S subunit (bS1-bS21) are shown. Target peptides are shown with underlined bold characters.

## Notes

### Competing Interest Statement

The authors have declared no competing interest.

## References

[1] J.L. Hsu, S.Y. Huang, N.H. Chow, S.H. Chen, Stable-isotope dimethyl labeling for quantitative proteomics, Anal. Chem. 75 (2003) 6843–6852, https://doi.org/10.1021/ac0348625.

[2] A. Thompson, J. Schäfer, K. Kuhn, S. Kienle, J. Schwarz, G. Schmidt, T. Neumann, R. Johnstone, A.K. Mohammed, C. Hamon, Tandem mass tags: a novel quantification strategy for comparative analysis of complex protein mixtures by MS/MS, Anal. Chem. 75 (2003) 1895–1904, https://doi.org/10.1021/ac0262560.

[3] P.L. Ross, Y.N. Huang, J.N. Marchese, B. Williamson, K. Parker, S. Hattan, N. Khainovski, S. Pillai, S. Dey, S. Daniels, S. Purkayastha, P. Juhasz, S. Martin, M. Bartlet-Jones, F. He, A. Jacobson, D.J. Pappin, Multiplexed protein quantitation in Saccharomyces cerevisiae using amine-reactive isobaric tagging reagents, Mol. Cell. Proteomics 3 (2004) 1154–1169, https://doi.org/10.1074/mcp.M400129-MCP200.

[4] L.V. DeSouza, A.M. Taylor, W. Li, M.S. Minkoff, A.D. Romaschin, T.J. Colgan, K.W. Siu, Multiple reaction monitoring of mTRAQ-labeled peptides enables absolute quantification of endogenous levels of a potential cancer marker in cancerous and normal endometrial tissues, J. Proteome Res. 7 (2008) 3525–3534, https://doi.org/10.1021/pr800312m.

[5] L. Dayon, M. Affolter, Progress and pitfalls of using isobaric mass tags for proteome profiling, Expert Rev. Proteomics 17 (2020) 149–161, https://doi.org/10.1080/14789450.2020.1731309.

[6] S.A. Gerber, J. Rush, O. Stemman, M.W. Kirschner, S.P. Gygi, Absolute quantification of proteins and phosphoproteins from cell lysates by tandem MS, Proc. Natl. Acad. Sci. U. S. A. 100 (2003) 6940–6945, https://doi.org/10.1073/pnas.0832254100.

[7] S.E. Ong, B. Blagoev, I. Kratchmarova, D.B. Kristensen, H. Steen, A. Pandey, M. Mann, Stable isotope labeling by amino acids in cell culture, SILAC, as a simple and accurate approach to expression proteomics, Mol. Cell. Proteomics 1 (2002) 376–386, https://doi.org/10.1074/mcp.m200025-mcp200.

[8] S.E. Ong, The expanding field of SILAC, Anal. Bioanal. Chem. 404 (2012) 967–976, https://doi.org/10.1007/s00216-012-5998-3.

[9] R. Narumi, K. Masuda, T. Tomonaga, J. Adachi, H.R. Ueda, Y. Shimizu, Cell-free synthesis of stable isotope-labeled internal standards for targeted quantitative proteomics, Synth. Syst. Biotechnol. 3 (2018) 97–104, https://doi.org/10.1016/j.synbio.2018.02.004.

[10] P.C. Zamecnik, E.B. Keller, Relation between phosphate energy donors and incorporation of labeled amino acids into proteins, J. Biol. Chem. 209 (1954) 337–354, https://doi.org/10.1016/S0021-9258(18)65561-9.

[11] A. Zemella, L. Thoring, C. Hoffmeister, S. Kubick, Cell-Free Protein Synthesis: Pros and Cons of Prokaryotic and Eukaryotic Systems, Chembiochem 16 (2015) 2420–2431, https://doi.org/10.1002/cbic.201500340.

[12] Y. Shimizu, A. Inoue, Y. Tomari, T. Suzuki, T. Yokogawa, K. Nishikawa, T. Ueda, Cell-free translation reconstituted with purified components, Nat. Biotechnol. 19 (2001) 751–755, https://doi.org/10.1038/90802.

[13] J. Yokoyama, T. Matsuda, S. Koshiba, N. Tochio, T. Kigawa, A practical method for cell-free protein synthesis to avoid stable isotope scrambling and dilution, Anal. Biochem. 411 (2011) 223–229, https://doi.org/10.1016/j.ab.2011.01.017.

[14] R. Narumi, Y. Shimizu, M. Ukai-Tadenuma, K.L. Ode, G.N. Kanda, Y. Shinohara, A. Sato, K. Matsumoto, H.R. Ueda, Mass spectrometry-based absolute quantification reveals rhythmic variation of mouse circadian clock proteins, Proc. Natl. Acad. Sci. U. S. A. 113 (2016) E3461–E3467, https://doi.org/10.1073/pnas.1603799113.

[15] J. Kamiie, S. Ohtsuki, R. Iwase, K. Ohmine, Y. Katsukura, K. Yanai, Y. Sekine, Y. Uchida, S. Ito, T. Terasaki, Quantitative atlas of membrane transporter proteins: development and application of a highly sensitive simultaneous LC/MS/MS method combined with novel in-silico peptide selection criteria, Pharm. Res. 25 (2008) 1469–1483, https://doi.org/10.1007/s11095-008-9532-4.

[16] R. Fields, The rapid determination of amino groups with TNBS, Methods Enzymol. 25 (1972) 464–468, https://doi.org/10.1016/S0076-6879(72)25042-X.

[17] J.H. Davis, J.R. Williamson, Structure and dynamics of bacterial ribosome biogenesis, Philos. Trans. R Soc. Lond. B Biol. Sci. 372 (2017) 20160181, https://doi.org/10.1098/rstb.2016.0181.

[18] M. Shimojo, K. Amikura, K. Masuda, T. Kanamori, T. Ueda, Y. Shimizu, In vitro reconstitution of functional small ribosomal subunit assembly for comprehensive analysis of ribosomal elements in E. coli, Commun. Biol. 3 (2020) 142, https://doi.org/10.1093/bioinformatics/btq054.

[19] D.M. Walther, P. Kasturi, M. Zheng, S. Pinkert, G. Vecchi, P. Ciryam, R.I. Morimoto, C.M. Dobson, M. Vendruscolo, M. Mann, F.U. Hartl, Widespread Proteome Remodeling and Aggregation in Aging C. elegans, Cell 161 (2015) 919–932, https://doi.org/10.1016/j.cell.2015.03.032.

[20] N. Slavov, S. Semrau, E. Airoldi, B. Budnik, A. van Oudenaarden, Differential Stoichiometry among Core Ribosomal Proteins, Cell Rep. 13 (2015) 865–873, https://doi.org/10.1016/j.celrep.2015.09.056.

[21] E. Emmott, M. Jovanovic, N. Slavov, Ribosome Stoichiometry: From Form to Function, Trends Biochem. Sci. 44 (2019) 95–109, https://doi.org/10.1016/j.tibs.2018.10.009.

[22] P. Mallick, M. Schirle, S.S. Chen, M.R. Flory, H. Lee, D. Martin, J. Ranish, B. Raught, R. Schmitt, T. Werner, B. Kuster, R. Aebersold, Computational prediction of proteotypic peptides for quantitative proteomics, Nat. Biotechnol. 25 (2007) 125–131, https://doi.org/10.1038/nbt1275.

[23] J.D. Worboys, J. Sinclair, Y. Yuan, C. Jørgensen, Systematic evaluation of quantotypic peptides for targeted analysis of the human kinome, Nat. Methods 11 (2014) 1041–1044, https://doi.org/10.1038/nmeth.3072.

[24] Y. Mohammed, D. Domański, A.M. Jackson, D.S. Smith, A.M. Deelder, M. Palmblad, C.H. Borchers, PeptidePicker: a scientific workflow with web interface for selecting appropriate peptides for targeted proteomics experiments, J. Proteomics 106 (2014) 151–161, https://doi.org/10.1016/j.jprot.2014.04.018.

[25] Z. Gao, C. Chang, J. Yang, Y. Zhu, Y. Fu, AP3: An Advanced Proteotypic Peptide Predictor for Targeted Proteomics by Incorporating Peptide Digestibility, Anal. Chem. 91 (2019) 8705–8711, https://doi.org/10.1021/acs.analchem.9b02520.

[26] M. Kumar, S.R. Joseph, M. Augsburg, A. Bogdanova, D. Drechsel, N.L. Vastenhouw, F. Buchholz, M. Gentzel, A. Shevchenko, MS Western, a Method of Multiplexed Absolute Protein Quantification is a Practical Alternative to Western Blotting, Mol. Cell. Proteomics 17 (2018) 384–396, https://doi.org/10.1074/mcp.O117.067082.

[27] M.V. Ivanov, I.A. Tarasova, L.I. Levitsky, E.M. Solovyeva, M.L. Pridatchenko, A.A. Lobas, J.A. Bubis, M.V. Gorshkov, MS/MS-Free Protein Identification in Complex Mixtures Using Multiple Enzymes with Complementary Specificity, J. Proteome Res. 16 (2017) 3989–3999, https://doi.org/10.1021/acs.jproteome.7b00365.

[28] Y. Shimizu, T. Ueda, PURE technology, Methods Mol. Biol. 607 (2010) 11–21, https://doi.org/10.1007/978-1-60327-331-2_2.

[29] T. Masuda, M. Tomita, Y. Ishihama, Phase transfer surfactant-aided trypsin digestion for membrane proteome analysis., J Proteome Res 7 (2008) 731–740, https://doi.org/10.1021/pr700658q.

[30] J. Rappsilber, M. Mann, Y. Ishihama, Protocol for micro-purification, enrichment, pre-fractionation and storage of peptides for proteomics using StageTips, Nat. Protoc. 2 (2007) 1896–1906, https://doi.org/10.1038/nprot.2007.261.

[31] B. MacLean, D.M. Tomazela, N. Shulman, M. Chambers, G.L. Finney, B. Frewen, R. Kern, D.L. Tabb, D.C. Liebler, M.J. MacCoss, Skyline: an open source document editor for creating and analyzing targeted proteomics experiments, Bioinformatics 26 (2010) 966–968, https://doi.org/10.1093/bioinformatics/btq054.

