## Supplementary Data 1 for "Versatile and multiplexed mass spectrometry-based absolute quantification with cell-free-synthesized internal standard peptides"

**> sP|P62894|CYC_BOVIN Cytochrome C**

MGDVEK/GK/K/IFVQK/CAQCHTVEK/GGK/HK/**TGPNLHGLFGR**/K/TGQAPGFSYTDANK/NK/GITWGEETLMEYLENPK/K/**YIPGTK**/MIFAGIK/K/K/GER/EDLIAYLK/K/ATNE

**> sP|P00698|LYSC_CHICK Lysozyme C**

MR/SLLILVLCFLPLAALGK/VFGR/CELAAAMK/R/**HGLDNYR**/GYSLGNWVCAAK/**FESNFNTQATNR**/**NTDGSTDYGILQINSR**/WWCNDGR/TPGSR/NLCNIPCSALLSSDITASVNCAK/K/IVSDGNGMNAWVAWR/NR/CK/GTDVQAWIR/GCR/L

**> sP|P00330|ADH1_YEAST Alcohol dehydrogenase 1**

MSIPETQK/**GVIFYESHGK**/LEYK/DIPVPK/PK/**ANELLINVK**/YSGVCHTDLHAWHGDWPLPVK/LPLVGGHEGAGVVVGMGENVK/GWK/**IGDYAGIK**/WLNGSCMACEYCELGNESNCPHADLSGYTHDGSFQQYATADAVQAAHIPQGTDLAQVAPILCAGITVYK/ALK/SANLMAGHWVAISGAAGGLGSLAVQYAK/AMGYR/VLGIDGGEGK/EELFR/SIGGEVFIDFTK/EK/DIVGAVLK/ATDGGAHGVINVSVSEAAIEASTR/YVR/ANGTTVLVGMPAGAK/CCSDVFNQVVK/SISIVGSYVGNR/ADTR/**EALDFFAR**/GLVK/SPIK/**VVGLSTLPEIYEK**/MEK/GQIVGR/**YVVDTSK**

**> sP|P02769|ALBU_BOVIN Serum albumin**

MK/WVTFISLLLLFSSAYSR/GVFR/R/DTHK/SEIAHR/FK/DLGEEHFK/GLVLIAFSQYLQQCPFDEHVK/**LVNELTEFAK**/TCVADESHAGCEK/SLHTLFGDELCK/VASLR/ETYGDMADCCEK/QEPER/NECFLSHK/DDSPDLPK/LK/PDPNTLCDEFK/ADEK/K/FWGK/YLYEIAR/R/HPYFYAPELLYYANK/YNGVFQECCQAEDK/GACLLPK/IETMR/EK/VLASSAR/QR/LR/CASIQK/FGER/ALK/**AWSVAR**/LSQK/FPK/**AEFVEVTK**/LVTDLTK/VHK/ECCHGDLLECADDR/ADLAK/YICDNQDTISSK/LK/ECCDK/PLLEK/SHCIAEVEK/DAIPENLPPLTADFAEDK/DVCK/**NYQEAK**/DAFLGSFLYEYSR/R/HPEYAVSVLLR/LAK/EYEATLEECCAK/DDPHACYSTVFDK/LK/**HLVDEPQNLIK**/QNCDQFEK/**LGEYGFQNALIVR**/YTR/K/VPQVSTPTLVEVSR/SLGK/VGTR/CCTK/PESER/MPCTEDYLSLILNR/LCVLHEK/TPVSEK/VTK/CCTESLVNR/R/PCFSALTPDETYVPK/AFDEK/LFTFHADICTLPDTEK/QIK/K/QTALVELLK/HK/PK/**ATEEQLK**/TVMENFVAFVDK/CCAADDK/EACFAVEGPK/LVVSTQTALA

**> sP|Q29443|TRFE_BOVIN Serotransferrin**

MR/PAVR/ALLACAVLGLCLADPER/TVR/WCTISTHEANK/CASFR/ENVLR/ILESGPFVSCVK/K/TSHMDCIK/AISNNEADAVTLDGGLVYEAGLK/PNNLK/PVVAEFHGTK/DNPQTHYYAVAVVK/K/DTDFK/LNELR/GK/K/SCHTGLGR/SAGWNIPMAK/LYK/**ELPDPQESIQR**/AAANFFSASCVPCADQSSFPK/LCQLCAGK/GTDK/CACSNHEPYFGYSGAFK/CLMEGAGDVAFVK/HSTVFDNLPNPEDR/K/NYELLCGDNTR/K/SVDDYQECYLAMVPSHAVVAR/TVGGK/EDVIWELLNHAQEHFGK/DK/PDNFQLFQSPHGK/DLLFK/DSADGFLK/IPSK/MDFELYLGYEYVTALQNLR/ESK/PPDSSK/DECMVK/WCAIGHQER/TK/CDR/WSGFSGGAIECETAENTEECIAK/IMK/GEADAMSLDGGYLYIAGK/CGLVPVLAENYK/TEGESCK/NTPEK/**GYLAVAVVK**/TSDANINWNNLK/DK/K/SCHTAVDR/TAGWNIPMGLLYSK/INNCK/FDEFFSAGCAPGSPR/NSSLCALCIGSEK/GTGK/ECVPNSNER/**YYGYTGAFR**/CLVEK/**GDVAFVK**/DQTVIQNTDGNNNEAWAK/NLK/K/ENFEVLCK/DGTR/K/PVTDAENCHLAR/GPNHAVVSR/K/DK/ATCVEK/ILNK/**QQDDFGK**/SVTDCTSNFCLFQSNSK/DLLFR/DDTK/CLASIAK/K/TYDSYLGDDYVR/AMTNLR/QCSTSK/LLEACTFHK/P

**> sP|P00722|BGAL_ECOLI Beta-galactosidase**

MTMITDSLAVVLQR/R/DWENPGVTQLNR/LAAHPPFASWR/NSEEAR/TDR/PSQQLR/SLNGEWR/FAWFPAPEAVPESWLECDLPEADTVVVPSNWQMHGYDAPIYTNVTYPITVNPPFVPTENPTGCYSLTFNVDESWLQEGQTR/IIFDGVNSAFHLWCNGR/**WVGYGQDSR**/LPSEFDLSAFLR/AGENR/LAVMVLR/WSDGSYLEDQDMWR/MSGIFR/DVSLLHK/PTTQISDFHVATR/**FNDDFSR**/AVLEAEVQMCGELR/DYLR/VTVSLWQGETQVASGTAPFGGEIIDER/**GGYADR**/VTLR/LNVENPK/**LWSAEIPNLYR**/AVVELHTADGTLIEAEACDVGFR/EVR/IENGLLLLNGK/PLLIR/GVNR/HEHHPLHGQVMDEQTMVQDILLMK/**QNNFNAVR**/CSHYPNHPLWYTLCDR/YGLYVVDEANIETHGMVPMNR/LTDDPR/WLPAMSER/VTR/MVQR/DR/NHPSVIIWSLGNESGHGANHDALYR/WIK/SVDPSR/PVQYEGGGADTTATDIICPMYAR/**VDEDQPFPAVPK**/WSIK/K/WLSLPGETR/PLILCEYAHAMGNSLGGFAK/YWQAFR/QYPR/LQGGFVWDWVDQSLIK/YDENGNPWSAYGGDFGDTPNDR/QFCMNGLVFADR/TPHPALTEAK/**HQQQFFQFR**/**LSGQTIEVTSEYLFR**/HSDNELLHWMVALDGK/PLASGEVPLDVAPQGK/QLIELPELPQPESAGQLWLTVR/VVQPNATAWSEAGHISAWQQWR/LAENLSVTLPAASHAIPHLTTSEMDFCIELGNK/R/WQFNR/QSGFLSQMWIGDK/K/QLLTPLR/DQFTR/APLDNDIGVSEATR/IDPNAWVER/WK/AAGHYQAEAALLQCTADTLADAVLITTAHAWQHQGK/TLFISR/K/TYR/IDGSGQMAITVDVEVASDTPHPAR/IGLNCQLAQVAER/**VNWLGLGPQENYPDR**/LTAACFDR/WDLPLSDMYTPYVFPSENGLR/CGTR/**ELNYGPHQWR**/**GDFQFNISR**/YSQQQLMETSHR/HLLHAEEGTWLNIDGFHMGIGGDDSWSPSVSAEFQLSAGR/YHYQLVWCQK
