## Supplementary Data 2 for "Versatile and multiplexed mass spectrometry-based absolute quantification with cell-free-synthesized internal standard peptides"

**>bS1**

MTESFAQLFEESLK/EIETR/PGSIVR/**GVVVAIDK**/DVVLVDAGLK/SESAIPAEQFK/NAQGELEIQVGDEVDVALDAVEDGFGETLLSR/EK/AK/R/HEAWITLEK/AYEDAETVTGVINGK/VK/GGFTVELNGIR/AFLPGSLVDVR/PVR/DTLHLEGK/ELEFK/VIK/LDQK/R/NNVVVSR/R/AVIESENSAER/DQLLENLQEGMEVK/GIVK/NLTDYGAFVDLGGVDGLLHITDMAWK/R/VK/**HPSEIVNVGDEITVK**/VLK/FDR/ER/TR/VSLGLK/QLGEDPWVAIAK/R/YPEGTK/LTGR/VTNLTDYGCFVEIEEGVEGLVHVSEMDWTNK/NIHPSK/VVNVGDVVEVMVLDIDEER/R/R/ISLGLK/QCK/ANPWQQFAETHNK/GDR/VEGK/IK/SITDFGIFIGLDGGIDGLVHLSDISWNVAGEEAVR/EYK/K/GDEIAAVVLQVDAER/ER/ISLGVK/QLAEDPFNNWVALNK/K/GAIVTGK/VTAVDAK/**GATVELADGVEGYLR**/ASEASR/DR/VEDATLVLSVGDEVEAK/FTGVDR/K/NR/AISLSVR/AK/DEADEK/DAIATVNK/QEDANFSNNAMAEAFK/AAK/GE

**>uS2**

MATVSMR/DMLK/**AGVHFGHQTR**/YWNPK/MK/PFIFGAR/NK/**VHIINLEK**/TVPMFNEALAELNK/IASR/K/GK/ILFVGTK/R/AASEAVK/DAALSCDQFFVNHR/WLGGMLTNWK/TVR/QSIK/R/LK/**DLETQSQDGTFDK**/LTK/K/EALMR/TR/ELEK/**LENSLGGIK**/DMGGLPDALFVIDADHEHIAIK/EANNLGIPVFAIVDTNSDPDGVDFVIPGNDDAIR/AVTLYLGAVAATVR/EGR/SQDLASQAEESFVEAE

**>uS3**

MGQK/**VHPNGIR**/LGIVK/PWNSTWFANTK/**EFADNLDSDFK**/VR/QYLTK/ELAK/ASVSR/IVIER/PAK/SIR/VTIHTAR/PGIVIGK/K/GEDVEK/LR/K/VVADIAGVPAQINIAEVR/K/PELDAK/LVADSITSQLER/R/VMFR/R/AMK/R/AVQNAMR/LGAK/GIK/VEVSGR/**LGGAEIAR**/TEWYR/EGR/**VPLHTLR**/ADIDYNTSEAHTTYGVIGVK/VWIFK/GEILGGMAAVEQPEK/PAAQPK/K/QQR/K/GR/K/

**>uS4**

MAR/YLGPK/LK/LSR/R/EGTDLFLK/SGVR/AIDTK/CK/IEQAPGQHGAR/K/PR/**LSDYGVQLR**/EK/QK/VR/R/IYGVLER/QFR/NYYK/EAAR/LK/**GNTGENLLALLEGR**/**LDNVVYR**/MGFGATR/AEAR/QLVSHK/AIMVNGR/VVNIASYQVSPNDVVSIR/EK/AK/K/QSR/VK/**AALELAEQR**/EK/PTWLEVDAGK/MEGTFK/R/K/PER/SDLSADINEHLIVELYSK/

**>uS5**

MAHIEK/QAGELQEK/LIAVNR/VSK/TVK/GGR/IFSFTALTVVGDGNGR/VGFGYGK/AR/EVPAAIQK/AMEK/AR/R/NMINVALNNGTLQHPVK/**GVHTGSR**/VFMQPASEGTGIIAGGAMR/**AVLEVAGVHNVLAK**/AYGSTNPINVVR/ATIDGLENMNSPEMVAAK/R/GK/**SVEEILGK**/

**>bS6**

MR/HYEIVFMVHPDQSEQVPGMIER/**YTAAITGAEGK**/IHR/**LEDWGR**/R/QLAYPINK/LHK/AHYVLMNVEAPQEVIDELETTFR/**FNDAVIR**/SMVMR/TK/HAVTEASPMVK/AK/DER/R/ER/R/DDFANETADDAEAGDSEEEEEE

**>uS7**

MPR/R/R/VIGQR/K/ILPDPK/FGSELLAK/FVNILMVDGK/K/STAESIVYSALETLAQR/SGK/SELEAFEVALENVR/PTVEVK/SR/R/VGGSTYQVPVEVR/PVR/R/NALAMR/WIVEAAR/K/R/GDK/SMALR/**LANELSDAAENK**/GTAVK/K/R/EDVHR/MAEANK/**AFAHYR**/WLSLR/**SFSHQAGASSK**/QPALGYLN

**>uS8**

MSMQDPIADMLTR/IR/NGQAANK/AAVTMPSSK/LK/**VAIANVLK**/**EEGFIEDFK**/VEGDTK/PELELTLK/YFQGK/**AVVESIQR**/VSR/PGLR/IYK/R/K/DELPK/VMAGLGIAVVSTSK/GVMTDR/AAR/QAGLGGEIICYVA

**>uS9**

MAENQYYGTGR/R/K/SSAAR/VFIK/PGNGK/**IVINQR**/**SLEQYFGR**/ETAR/MVVR/QPLELVDMVEK/**LDLYITVK**/GGGISGQAGAIR/HGITR/ALMEYDESLR/SELR/K/AGFVTR/DAR/QVER/K/K/VGLR/K/AR/R/R/PQFSK/R/

**>uS10**

MQNQR/IR/IR/LK/AFDHR/LIDQATAEIVETAK/R/TGAQVR/GPIPLPTR/K/ER/**FTVLISPHVNK**/DAR/**DQYEIR**/THLR/**LVDIVEPTEK**/TVDALMR/LDLAAGVDVQISLG

**>uS11**

MAK/APIR/AR/K/R/VR/K/QVSDGVAHIHASFNNTIVTITDR/QGNALGWATAGGSGFR/GSR/K/**STPFAAQVAAER**/CADAVK/EYGIK/NLEVMVK/**GPGPGR**/ESTIR/**ALNAAGFR**/ITNITDVTPIPHNGCR/PPK/K/R/R/V

**>uS12**

MATVNQLVR/K/PR/AR/K/VAK/**SNVPALEACPQK**/R/GVCTR/**VYTTTPK**/K/PNSALR/K/VCR/VR/LTNGFEVTSYIGGEGHNLQEHSVILIR/GGR/VK/DLPGVR/YHTVR/**GALDCSGVK**/DR/K/QAR/SK/YGVK/R/PK/A

**>uS13**

MAR/**IAGINIPDHK**/**HAVIALTSIYGVGK**/TR/SK/**AILAAAGIAEDVK**/**ISELSEGQIDTLR**/DEVAK/**FVVEGDLR**/R/EISMSIK/R/LMDLGCYR/GLR/HR/R/GLPVR/GQR/TK/TNAR/TR/K/GPR/K/PIK/K/

**>uS14**

MAK/QSMK/AR/EVK/R/VALADK/YFAK/R/AELK/**AIISDVNASDEDR**/**WNAVLK**/**LQTLPR**/DSSPSR/QR/NR/CR/QTGR/PHGFLR/K/FGLSR/IK/VR/EAAMR/**GEIPGLK**/K/ASW

**>uS15**

MSLSTEATAK/**IVSEFGR**/DANDTGSTEVQVALLTAQINHLQGHFAEHK/K/DHHSR/R/GLLR/MVSQR/R/K/**LLDYLK**/R/K/DVAR/**YTQLIER**/LGLR/R/

**>bS16**

MVTIR/LAR/HGAK/K/R/PFYQVVVADSR/NAR/NGR/FIER/**VGFFNPIASEK**/EEGTR/LDLDR/**IAHWVGQGATISDR**/**VAALIK**/EVNK/AA

**>uS17**

MTDK/IR/TLQGR/VVSDK/MEK/**SIVVAIER**/FVK/**HPIYGK**/FIK/R/TTK/**LHVHDENNECGIGDVVEIR**/ECR/PLSK/TK/**SWTLVR**/VVEK/AVL

**>bS18**

MAR/YFR/R/R/K/FCR/**FTAEGVQEIDYK**/DIATLK/**NYITESGK**/IVPSR/ITGTR/AK/YQR/QLAR/AIK/R/AR/**YLSLLPYTDR**/HQ

**>uS19**

MPR/SLK/K/**GPFIDLHLLK**/K/VEK/AVESGDK/K/PLR/TWSR/R/STIFPNMIGLTIAVHNGR/QHVPVFVTDEMVGHK/**LGEFAPTR**/TYR/GHAADK/K/AK/K/K/

**>bS20**

MANIK/SAK/K/R/AIQSEK/AR/K/HNASR/R/SMMR/TFIK/K/**VYAAIEAGDK**/AAAQK/AFNEMQPIVDR/QAAK/**GLIHK**/NK/AAR/HK/**ANLTAQINK**/LA

**>bS21**

MPVIK/VR/**ENEPFDVALR**/R/FK/R/SCEK/**AGVLAEVR**/R/R/EFYEK/PTTER/K/R/AK/ASAVK/R/HAK/K/LAR/ENAR/R/TR/LY
